# Cisplatin-induced oxidative stress regulates YAP to modulate epigenome promoting survival of osteosarcoma cells

**DOI:** 10.1101/2025.08.25.672065

**Authors:** Ankita Daiya, Chinmay Nayak, Rajdeep Chowdhury, Shibasish Chowdhury, Sudeshna Mukherjee

**Affiliations:** Department of Biological Sciences, Birla Institute of Technology & Science, Pilani, Pilani Campus, Rajasthan, India 333031

**Keywords:** Cisplatin, Osteosarcoma, ROS, YAP, EZH2, H3K27me3

## Abstract

The widely used chemotherapeutic drug cisplatin (CDDP) is an integral part of the pre-operative chemotherapy protocol for high-grade osteosarcoma (OS). However, despite an aggressive treatment regimen, drug refractoriness is a major hindrance to successful therapy. We previously identified key transcriptomic alterations essential for the survival of OS cells following CDDP exposure. In the present study, we further demonstrate that CDDP treatment resulted in a ROS-dependent enrichment of the repressive histone mark H3K27me3 at the upstream promoter regions of growth-promoting genes such as CCNA2, and on the promoter of the negative regulator of Yes-Associated Protein (YAP)-LATS1, thereby contributing to their transcriptional repression. This was associated with a growth arrest, and quenching of ROS with N-acetyl cysteine (NAC) reversed it. Importantly, repression of LATS1 led to an increased nuclear localization of YAP, while pharmacological or genetic ablation of YAP reduced CDDP-mediated induction of repressive marks. YAP was further found to co-localize and co-immunoprecipitate with the Polycomb Repressive Complex 2 (PRC2) catalytic member-the histone methyl transferase-EZH2, indicating its putative role in mediating transcriptional repression. In lieu of the above, inhibition of YAP or reversal of the repressive chromatin state using a histone deacetylase (HDAC) inhibitor sensitized OS cells to a low-dose CDDP treatment as well. Overall, the present study demonstrates an interplay between oxidative stress, epigenetics, and YAP in modulating OS cell fate post CDDP exposure.

## 1. Introduction

Osteosarcoma (OS) is one of the prevalent types of primary malignant bone tumors and is clinically aggressive (1, 2). Even with advances in intensive medical research, 40% of OS patients do succumb to the disease. Given that the majority of incidents occur in young people under the age of 20, this malignancy is considered rare, and therefore, there is a scarcity of reports examining the molecular biology of the disease (3). Cisplatin (CDDP) is a primary line of chemotherapeutic drugs that is often used in the treatment of OS (4). It is known to cause DNA damage leading to oxidative stress, which can trigger a multitude of responses leading to genomic instability, structural changes, and as well as epigenetic changes (5, 6). For instance, in some cancers, 8-hydroxydeoxyguanosine (8-OHdG) levels are elevated under high oxidative stress conditions that induce conformational change, transforming active chromatin state into a repressive one; this change affects the methylation patterns of major tumor suppressor genes suggesting that drug treatment, oxidative stress, and epigenome have intricate links (7, 8, 9). However, as extensively as the DNA methylation states are studied, histone modulation by reactive oxygen species (ROS), under drug stress, is still poorly explored.

Herein, identifying the signaling pathways acting as the connecting link between oxidative stress and epigenomic modifications under drug stress can provide further hints on novel targets. In this regard, the Hippo pathway is majorly involved in cellular proliferation and death during organ development (10). Yes-associated protein (YAP), a downstream effector of this pathway, is a transcriptional coactivator that undergoes a cycle of phosphorylation or dephosphorylation. Disruption of Hippo signaling leads to reduced phosphorylation of YAP, facilitating its translocation to the nucleus. Within the nucleus, YAP regulates key cellular processes, including proliferation, survival, and growth. Increasing evidence now indicates that elevated expression of YAP is associated with poor prognosis across multiple cancer types. (11–13). However, the role of YAP in regulating oxidative stress response and epigenetic changes is not completely understood. For pediatric bone cancers like OS, YAP is established to be involved in more than one way-starting with angiogenesis to metastatic dissemination, which underlines the importance of studying YAP in the context of epigenomic changes (14, 15).

Earlier studies demonstrate that oxidative stress can lead to dysregulation in Hippo/YAP pathway components and *vice versa*. In NSCLCs, oxidative stress causes miR-25 to be overexpressed, which in turn inhibits LATS1, a negative regulator of YAP, thereby rendering YAP active (16). Similarly, YAP, when activated, is reported to form a complex with FOXO1 and occupy the promoter of antioxidant genes such as catalase (17). Similarly, in cancers such as angiosarcoma, endothelial cells with enhanced YAP activity are more resistant to oxidative stress (18). These studies suggest that ROS might have a role in regulating the Hippo/YAP pathway to promote chemoresistance. Interestingly, YAP, a critical regulator of developmental programs, has also been implicated in modulating chromatin architecture, including interactions with chromatin remodeling complexes such as NuRD (19, 20). However, the precise mechanisms underlying the crosstalk between YAP and epigenetic modifiers remain to be fully elucidated.

In the current study, we exhibit that a ‘sub-lethal’ dose of CDDP can lead to a growth-arrest state but minimal cellular death. Additionally, we show that CDDP, even at lower doses, is potent enough to induce H3K27me3 alterations along with changes in the expression pattern of its catalyst, EZH2. Further, we explored the crosstalk between ROS, the Hippo/YAP pathway, and the regulation of the histone repressive marks. Deeper insights into the molecular mechanisms of YAP and EZH2 can be useful in designing appropriate therapeutic strategies against the recurrence of rarer but difficult-to-treat cancers like osteosarcoma.

## 2. Materials and Methods

### 2.1. Chemicals and reagents

Cisplatin (Cat. #232120-50M), 2′,7′-dichlorofluorescin diacetate (DCFDA, #D6883), propidium iodide (PI; #P4864), Bradford reagent (#B6916), verteporfin (#SML0534-5MG), TRI reagent® (#T9424), DAPI (#D9542), and RIPA buffer (#R0278) were procured from Sigma-Aldrich. N-acetyl-L-cysteine (NAC; 47866) and MTT reagent (#33611) were obtained from SRL. Suberoylanilide hydroxamic acid (SAHA; #H1388) was purchased from TCI, and the EZH2 inhibitor GSK-126 (#5005800001) from Merck. Chromatin immunoprecipitation was performed using the MAGnify™ Chromatin Immunoprecipitation System (#492024), and apoptosis assays were conducted using Annexin V-FITC Conjugate (#A13199), both from Thermo Fisher Scientific. Immunofluorescence specific secondary antibodies Anti-Rabbit IgG Alexa Fluor 555 (#A32732) and Anti-Mouse IgG Alexa Fluor 488 (#A32723) were obtained from Invitrogen. Mitochondrial Membrane Potential Probe (JC-1 dye #T3168), MitoSOX Red (#M36008), and Lipofectamine 3000 (#L3000001) were also sourced from Invitrogen. The Luciferase Assay System (#E1500) was purchased from Promega. Western blot detection reagents, including the Clarity Western ECL Substrates (#1705061), PCR iTaq™ Universal SYBR Green Supermix (#1725121), and iScript™ cDNA Synthesis Kit (#170-8891), were acquired from Bio-Rad. Primary antibodies against YAP (#sc-101199) and GAPDH (#sc-365062) were obtained from Santa Cruz Biotechnology. Additional primary antibodies were sourced from Cell Signaling Technology (CST): anti-H3K27me3 (#9733S), anti-EZH2 (#5246S), anti-YAP (#8418S), anti-LATS1 (#3477T), anti-CYR61 (#14479), anti-p21 (#2947S), and anti-PCNA (#13110S). The HRP-conjugated secondary antibodies, anti-mouse IgG (Cat. #7076S) and anti-rabbit IgG (Cat. #7074P2), were obtained from CST. siRNA targeting YAP (ID #107951) and scrambled control siRNA (ID #32-6976) were obtained from commercial sources. The 8XGTIIC luciferase reporter plasmid (Addgene plasmid #34615) was kindly provided by Dr. Stefano Piccolo. Minimum Essential Medium Eagle (MEM; Cat. #AL047S) was obtained from HiMedia.

### 2.2. Cell culture

The human osteosarcoma cell line HOS (CRL-1543) was obtained from the National Centre for Cell Science (NCCS), Pune, India. Cells were maintained in Minimal Essential Medium (MEM; HiMedia) supplemented with 10% fetal bovine serum (FBS; Gibco), under standard culture conditions at 37°C with 5% COL. For experiments, cells were cultured until 60–70% confluency, washed with phosphate-buffered saline (PBS), and subsequently reseeded in fresh medium before administration of the specified drug treatment(s).

### 2.3. Cell viability assay

Cell viability in response to varying concentrations of CDDP was assessed using the MTT assay. A total of 6 × 10³ cells per well were seeded into 96-well plates. After allowing the cells to adhere and attain characteristic morphology, they were treated with increasing concentrations of Cisplatin for 24, 48, or 72 hours. Following treatment, MTT reagent was added to each well and incubated for 3 hours to allow for the formation of formazan crystals. The resulting crystals were solubilized using dimethyl sulfoxide (DMSO), and absorbance was measured at 570 nm with a reference filter at 630 nm using a Multiskan GO microplate spectrophotometer (Thermo Scientific). Cell viability was calculated using the formula: % Cell Viability = (Mean absorbance of Cisplatin-treated cells / Mean absorbance of control cells) × 100 (44).

### 2.4. Estimation of intracellular reactive oxygen species (ROS)

Reactive oxygen species levels were measured using DCFH-DA, and for that, 6×10^3^ cells per well were seeded. To inhibit basal ROS, NAC was added two hours before drug treatment. Following treatment with cisplatin, cells were washed with 1×PBS and incubated with 100LμL of 20 μM DCFDA for 30 minutes at 37L°C. Fluorescence was measured using a microplate reader at an excitation wavelength of 485 nm and emission wavelength of 530Lnm (44).

### 2.5. siRNA transfection

6×10^3^ cells were seeded for this experiment in a 6-well cell culture plate. Upon reaching around 70% confluency, cells were transfected with designated siRNAs (40 nM) using Lipofectamine 3000 reagent according to Invitrogen’s instruction leaflet. Scrambled siRNA was used as a negative control.

### 2.6. Immunoblotting

Treated cells were lysed using a modified radioimmunoprecipitation assay (RIPA) buffer. Total protein concentration was determined using the Bradford assay. For preparation of lysates, a 5× loading buffer was added, followed by denaturation at 100L°C for 10 minutes. Equal amounts of protein were resolved by SDS-PAGE and subsequently transferred onto polyvinylidene difluoride (PVDF) membranes. Membranes were blocked with either 5% skimmed milk or 3% bovine serum albumin (BSA) in TBST, depending on the antibody used. Before antibody incubation, blots were sectioned as appropriate. Then membranes were subsequently incubated with specific primary antibodies and, if required, stripped and re-incubated with additional primary antibodies, followed by appropriate HRP-conjugated secondary antibodies. Protein bands were visualized using an enhanced chemiluminescence (ECL) detection system, and densitometric analysis was performed using ImageJ software. (44).

### 2.7. Annexin V/PI apoptosis assay

Cells were seeded in 6 cm culture dishes at an appropriate density and treated with CDDP, Verteporfin (VP), GSK-126, or SAHA for 24 hours. Following treatment, cells were harvested and washed with 1× phosphate-buffered saline (PBS). Apoptosis analysis was performed as per a previously established protocol using the CytoFLEX flow cytometer (Beckman Coulter) (44). The acquired data were analyzed using CytExpert software. The percentages of early and late apoptotic cells were quantified, and results were represented as fold change in apoptotic cell populations using bar graphs.

### 2.8. Estimation of mitochondrial membrane potential

A total of 1 × 10L cells were seeded in 6 cm culture dishes and treated with Cisplatin for 24 hours. Following treatment, mitochondrial membrane potential was assessed using JC-1 dye at a final working concentration of 70 nM. Cells were incubated with the dye for 30 minutes at 37 °C, after which they were harvested and washed twice with phosphate-buffered saline (PBS). The cell pellet was resuspended in 500 μL of 1× PBS, and fluorescence data were acquired using a CytoFLEX flow cytometer (Beckman Coulter). Data analysis was performed using CytExpert software. Mitochondrial membrane depolarization was quantified as the ratio of red to green fluorescence and represented graphically as a bar chart

### 2.9. RNA isolation, cDNA synthesis, and qPCR

Total cellular RNA was extracted using TRI reagent^®^ and quantified. cDNA was synthesized using 2 ug of RNA. cDNA was amplified for the desired genes in the Quant Studio 3.0 Real-time PCR System. Beta Actin was taken as a housekeeping control. Pfaffl’s method was used to calculate relative expression (45). The primers used and their sequences are provided in the following table (**Table 1**).

**Table 1.**
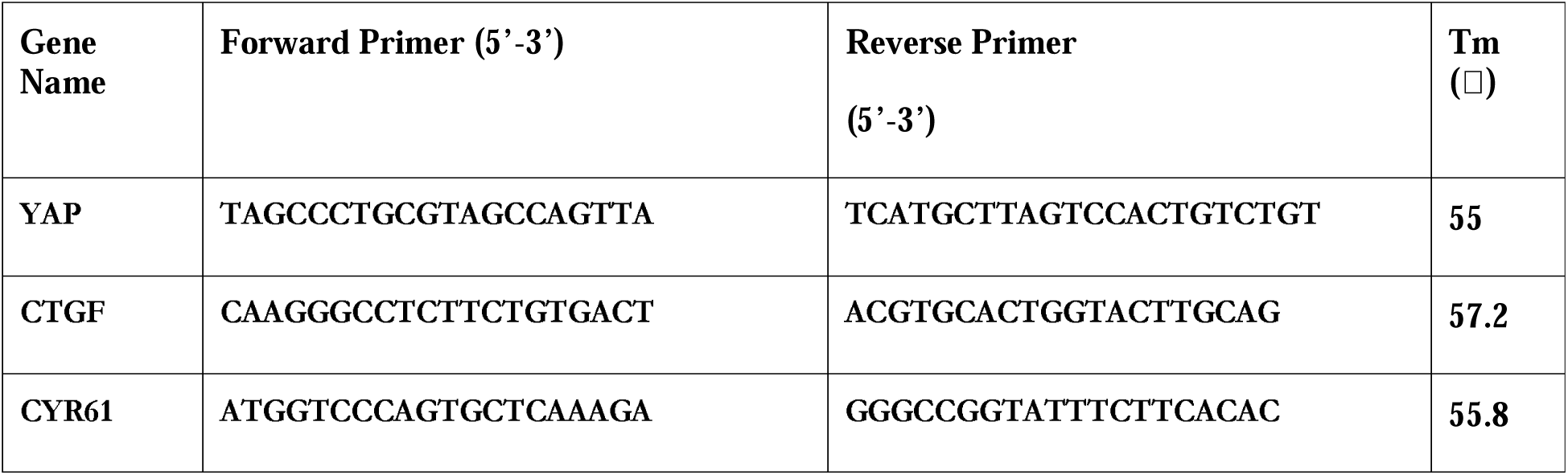
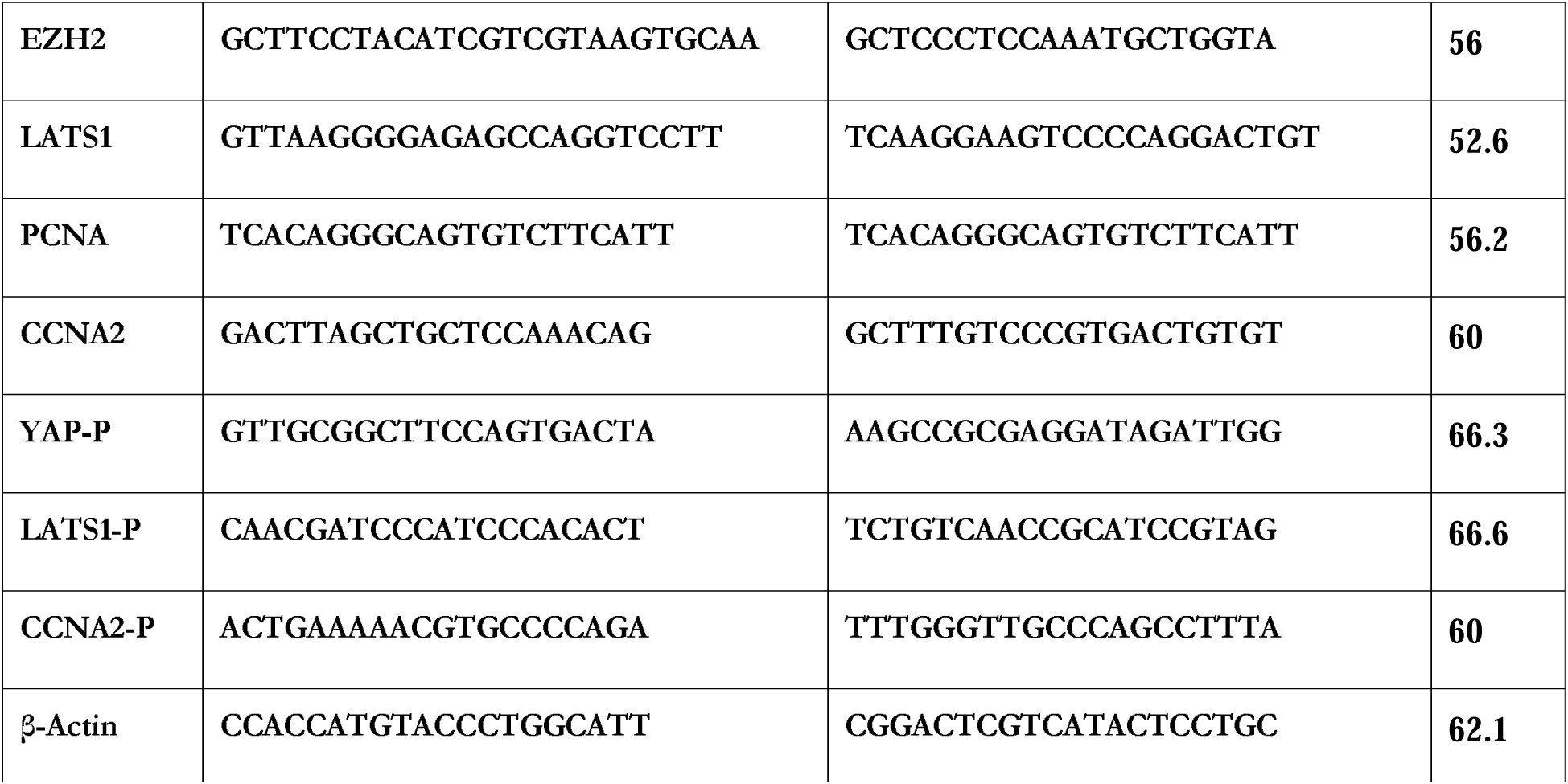
List of primers used in the study

### 2.10. Immunofluorescence

Cells were seeded at a density of 2.5 × 10L cells per well on sterile coverslips placed in 6-well plates. After reaching the required confluence, cells were exposed to the designated drug treatments. Following treatment, cells were washed thoroughly with 1× phosphate-buffered saline (PBS) and fixed with 2% paraformaldehyde (PFA) for 10 minutes at room temperature. After three washes with 1× PBS, cells were permeabilized with 0.1% Triton X-100 for 1 minute and subsequently blocked with 2.5% bovine serum albumin (BSA) for 60 minutes. Primary antibody incubation (1:1000 dilution in 2.5% BSA) was carried out overnight at 4L°C. The following day, after two washes with 1× PBS, cells were incubated with Alexa Fluor-conjugated secondary antibodies diluted in 1:2000 in 2.5% BSA for 60 minutes at room temperature. Nuclei were counterstained with DAPI for 10 minutes. After mounting coverslips onto glass slides with 70% glycerol, samples were visualized using the Zeiss Axio Observer.Z1/7 Apotome microscope. Image acquisition and analysis were performed using Zen 2.3 SP1 software by Zeiss.

### 2.11. Cell cycle analysis

A total of 1 × 10L cells were seeded in 6 cm culture dishes and subjected to the indicated drug treatments. Following exposure, cells were harvested, washed with 1× phosphate-buffered saline (PBS), and centrifuged at 5000 rpm for 10 minutes at 4 °C. The cell pellet was resuspended in ice-cold 70% ethanol while gently vortexing to ensure uniform fixation and stored overnight at 4 °C. The next day, cells were washed and resuspended in 500 μL of 1× PBS containing 4 μL of propidium iodide (PI), followed by incubation in the dark for 10 minutes at room temperature. Cellular DNA content was quantified based on PI fluorescence using the CytoFLEX flow cytometer (Beckman Coulter). Data acquisition and analysis were performed using CytExpert software.

### 2.12. ChIP-qPCR

ChIP assay was conducted utilizing the MAGnify Immunoprecipitation system (Invitrogen, Cat. #49-2024). OS cells were grown in 10 cm culture plates and treated with specified drug. After treatment, cells were harvested and rinsed three times with PBS. Cross-linking of protein-DNA complexes was achieved by incubating the cells with 1% formaldehyde, which was then neutralized by adding 1.25 M glycine. Following additional PBS washes, cells were lysed in buffer containing a protease inhibitor cocktail (50 µL per 10^6^ cells) and incubated on ice for 10 minutes with intermittent vortexing. Chromatin fragmentation was performed through sonication for 60 cycles (45 seconds on, 15 seconds off), and the efficiency of shearing was confirmed by electrophoresis on 1.2% agarose gel. The fragmented chromatin was incubated overnight with an anti-H3K27me3 antibody (dilution 1:50) to capture specific histone modifications. After a series of washes, immune complexes were eluted and reverse cross-linked. DNA purification was completed using proteinase K digestion. The enrichment of targeted DNA sequences from promoters of LATS1, YAP, and CCNA2 genes was quantified by ChIP-qPCR using specific primers.

### 2.13. Luciferase assay

Cells were seeded at a density of 2.5 × 10L cells per well in 6-well plates and transfected with the 8×GTIIC-luciferase reporter plasmid for YAP activity using Lipofectamine reagent (as instructed by the manufacturer). Following 4 to 6 hours of incubation post-transfection, cells were treated with the indicated drugs. Luciferase activity was assessed using Promega’s Luciferase Assay Kit according to the manufacturer’s protocol. Briefly, cells were washed with 1× PBS and lysed using the lysis buffer provided in the kit. Lysates were collected by scraping and centrifuged at 2500 rpm for 15 minutes. Equal volumes of clarified lysate were mixed with 100 μL of Luciferase Assay Reagent, and luminescence was immediately measured using the GloMax® 20/20 luminometer (Promega)

### 2.14. Statistical analysis

Statistical analyses were performed using GraphPad Prism software version 8.0. Depending on the experimental design, either an unpaired two-tailed Student’s t-test or one-way analysis of variance (ANOVA) was applied to assess the significance of differences between treatment groups and controls. For multiple comparisons, Tukey’s test was employed. p-values are indicated on individual graphs. Statistical significance was defined as p ≤ 0.05 and represented as follows: * for p ≤ 0.05, ** for p ≤ 0.01, and *** for p ≤ 0.001.

## 3. Results

### 3.1. Cisplatin induces a repressive chromatin mark in OS cells

Patient treatment regimens are now dependent on combinatorial treatment strategies in different cancers (21). This generally includes lower doses of multiple standard drugs to achieve an enhanced therapeutic effect, coupled with reduced off-target toxicity, compared to independent high doses of drug treatment. In this regard, while several studies delineate molecular alterations at IC50 or toxic doses of standard anti-cancer drugs, the effects of low-dose chemotherapeutics are rarely explored. Studies with low doses of standard drugs can ideally facilitate the identification of novel markers that can be simultaneously targeted along with the parent drug as a combinatorial treatment strategy for enhanced efficacy. Given the above, we initially treated human osteosarcoma (OS) cells with increasing doses of cisplatin (CDDP) and identified the IC50 of CDDP to be around 12 µM at 24h (**Fig 1A)**. Thereafter, a four times lower dose of CDDP (3 µM) was selected as a ‘sub-toxic’ dose and used for subsequent studies to identify putative targetable molecular markers which can be co-targeted. Herein, the combinatorial treatment strategies are often selected through the inclusion of drugs with different targets or mechanisms of action to have added therapeutic benefit. Therefore, since CDDP is known to target DNA, we were interested in exploring coupled epigenetic alterations associated with the drug. Interestingly, we observed that even a low dose of CDDP treatment can result in a significant increase in the repressive chromatin mark, as evidenced through increased transcript and protein levels of the gene-Enhancer of Zeste Homolog 2 (EZH2), a widely known methyl transferase (**Fig 1B-C**). In addition, EZH2 exhibited a pronounced shift toward nuclear localization following CDDP treatment **(Fig 1D)**. As above, the substrate of EZH2-H3K27me3 levels also showed a simultaneous increase and nuclear localization post CDDP exposure **(Fig 1E-F)**. A chromatin repression is often also associated with histone deacetylation (22). Accordingly, we observed an enhanced transcript level of several HDACs as shown in **Supplementary Fig 1A-C**. Overall, our results suggest that even at a sub-toxic low dose of the chemotherapeutic DNA-damaging drug -CDDP, there is associated chromatin repression, and therefore, epigenetic targeting can be further explored for putative enhanced efficacy.

**Figure 1.**
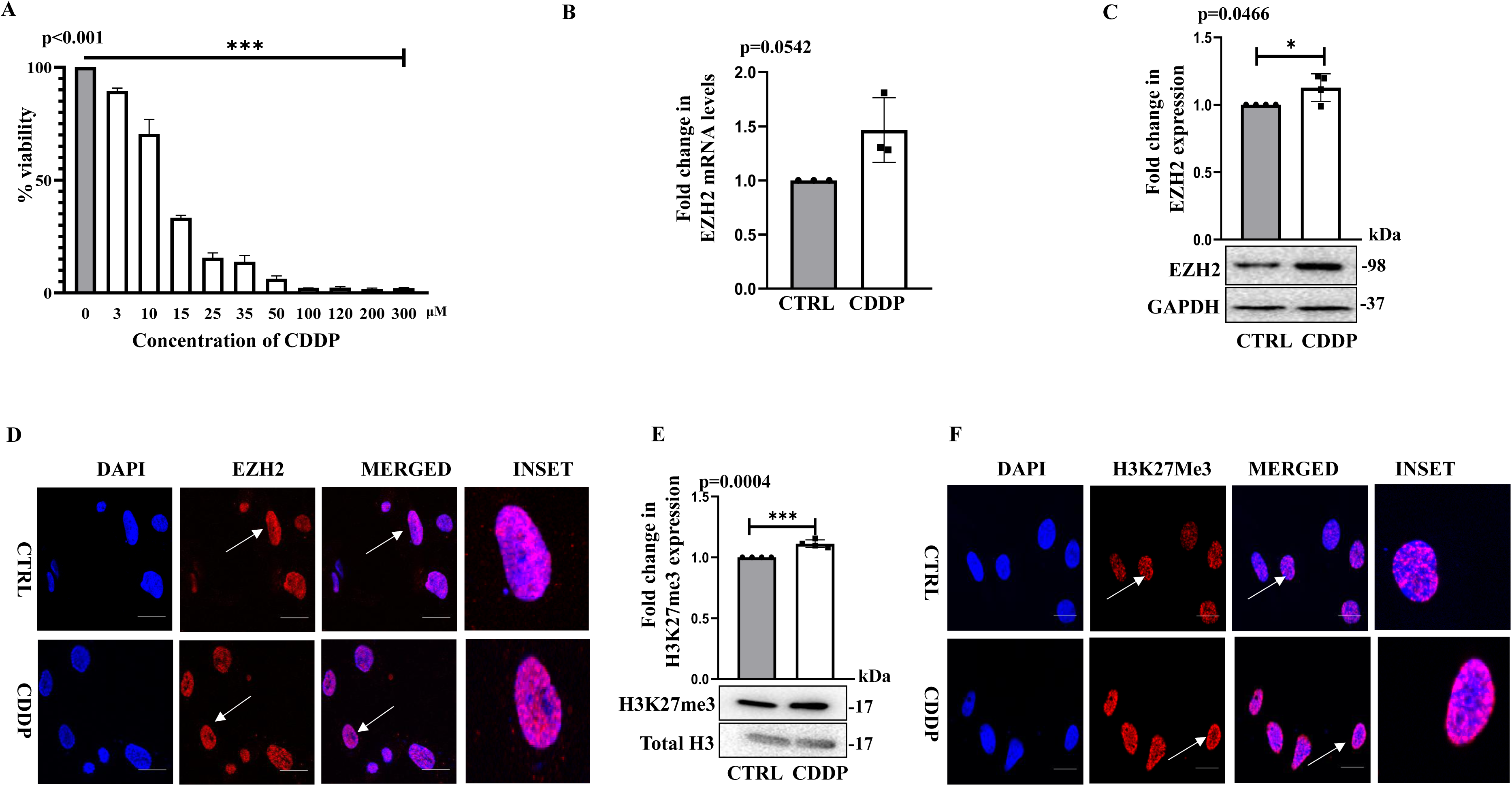
Cisplatin causes upregulation of repressive chromatin marks in HOS cells. **(A)** Cell Viability (MTT) assay at different concentrations of CDDP for 24h in OS cells. **(B)** Change in mRNA expression of EZH2 at 24h when exposed to a sub-lethal dose of CDDP (3 µM). **(C)** Immunoblot analysis showing expression of EZH2 at 24h; GAPDH served as the loading control. **(D)** Immunofluorescence staining showing the expression of EZH2 at 24h (Scale Bar: 20 μm). **(E)** Immunoblot analysis showing expression of H3K27me3 at 24h; total H3 served as the loading control. **(F)** Immunofluorescence staining showing the expression of H3K27me3 at 24h (Scale Bar: 20 μm). All values are represented as mean±SD; n=3. Unpaired t-test was applied with (*) p<0.05 to estimate the significance when two groups were compared. CTRL represents untreated cells.

### 3.2. Increase in repressive chromatin mark correlates with cellular growth arrest

Apart from an increase in histone repressive mark, resulting in putative transcriptional repression, the low dose of CDDP also induced a non-proliferative phenotype marked by an altered cell cycle distribution pattern **(Fig 2A**). In parallel, we also observed a reduced expression of cellular proliferation/replication-associated genes, like PCNA and CCNA2 (CyclinA2) **(Fig 2B and 2C)**. This was coupled to an increased expression of the cell cycle inhibitor-CDKN1A/p21-a cyclin-dependent kinase inhibitor, confirming the non-proliferative state of the cells (**Fig 2D)**. To elucidate the epigenetic mechanisms governing the non-proliferative cellular state, we assessed H3K27me3 enrichment at the upstream promoter region of CCNA2 via chromatin immunoprecipitation coupled with quantitative PCR (ChIP-qPCR). Importantly, we observed a significant enrichment of H3K27me3 on the probed CCNA2 promoter upstream element (**Fig 2E, Supplementary Figure 1 D**). Our data suggests that the drug-CDDP renders the OS cells in a non-dividing-like state, mediated by epigenetic mechanisms involving histone methylation at selective genetic loci.

**Figure 2.**
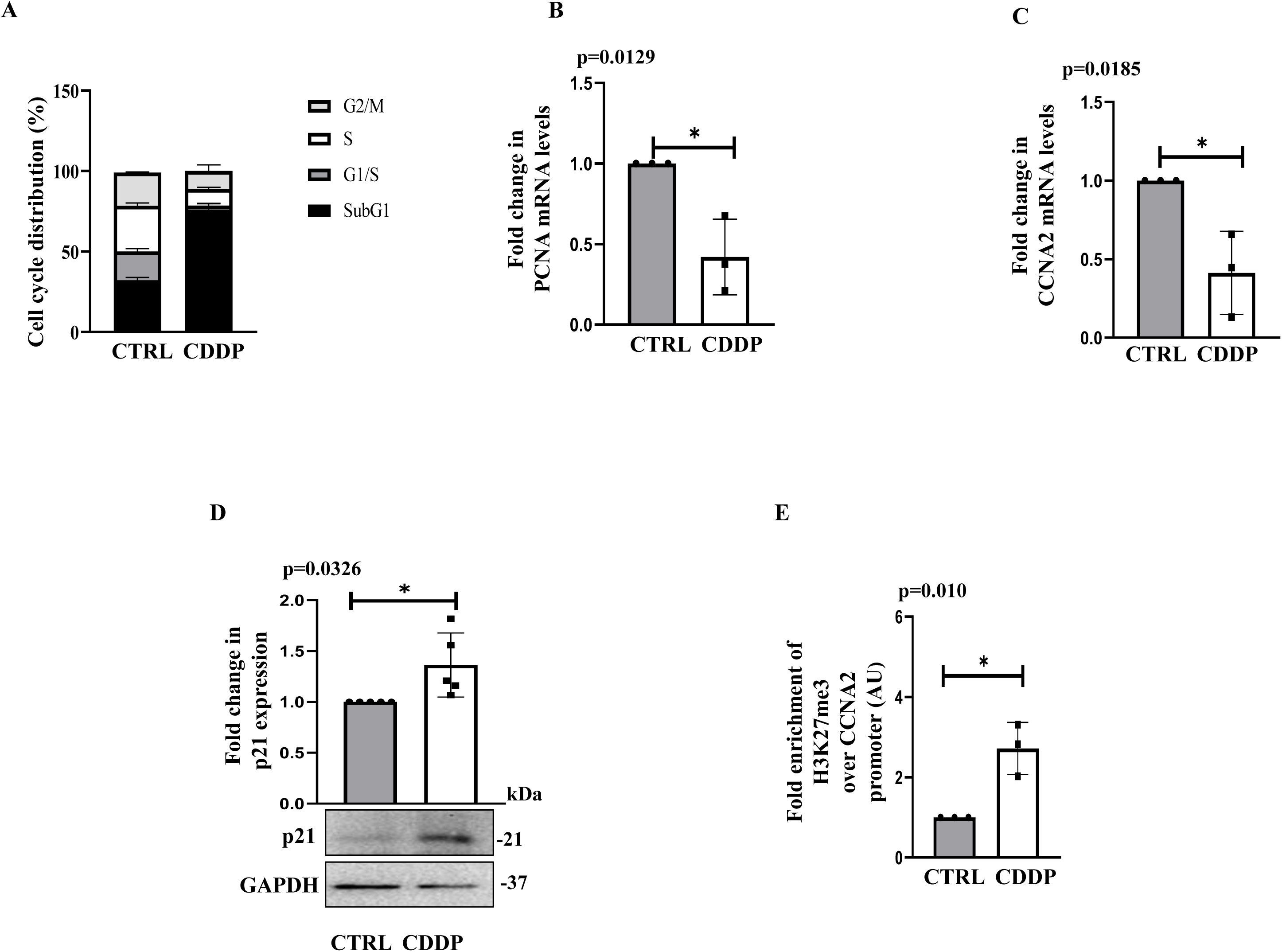
An increase in repressive chromatin marks is associated with growth arrest in HOS cells. **(A)** Cell cycle analysis through flow cytometry at 24h after CDDP exposure with respect to untreated HOS cells. **(B)** Change in mRNA expression of PCNA at 24h. **(C)** Change in mRNA expression of cell cycle gene CCNA2 at 24h. **(D)** Immunoblot analysis showing expression of p21 at 24h; GAPDH served as the loading control. **(E)** Enrichment of H3K27me3 over the cell cycle genes (CCNA2) as analysed through ChIP-qPCR. All values are represented as mean±SD; n=3. Unpaired t-test was applied with (*) p<0.05 to estimate the significance when two groups were compared. CTRL represents untreated cells.

### 3.3. Reactive Oxygen Species regulate induction of the repressive epigenomic mark

To understand the mediator of epigenetic alterations under drug stress, we assessed the generation of cellular reactive oxygen species (ROS), given that genotoxic agents such as anticancer drugs are frequently associated with elevated ROS levels. (23). Intracellular ROS levels were quantified using the DCFDA assay. As illustrated in **Fig 3A**, a significant increase in ROS was observed following CDDP treatment across all the examined time points. Considering that mitochondria are a major source of intracellular ROS, we also evaluated mitochondrial ROS levels and found them to be elevated post-CDDP exposure **(Fig 3B)**. This increase was associated with changes in mitochondrial membrane potential, as assessed by JC-1 staining **(Supplementary Figure 2A)**. Notably, a sub-toxic dose of CDDP, while sufficient to induce ROS, did not result in significant cell death (data not shown), instead exerting a predominantly cytostatic effect, as previously described. Given the role of oxidative stress in modulating epigenetic states, we next sought to understand whether ROS contributes to the modulation of H3K27me3 enrichment following CDDP treatment (5, 24).

**Figure 3.**
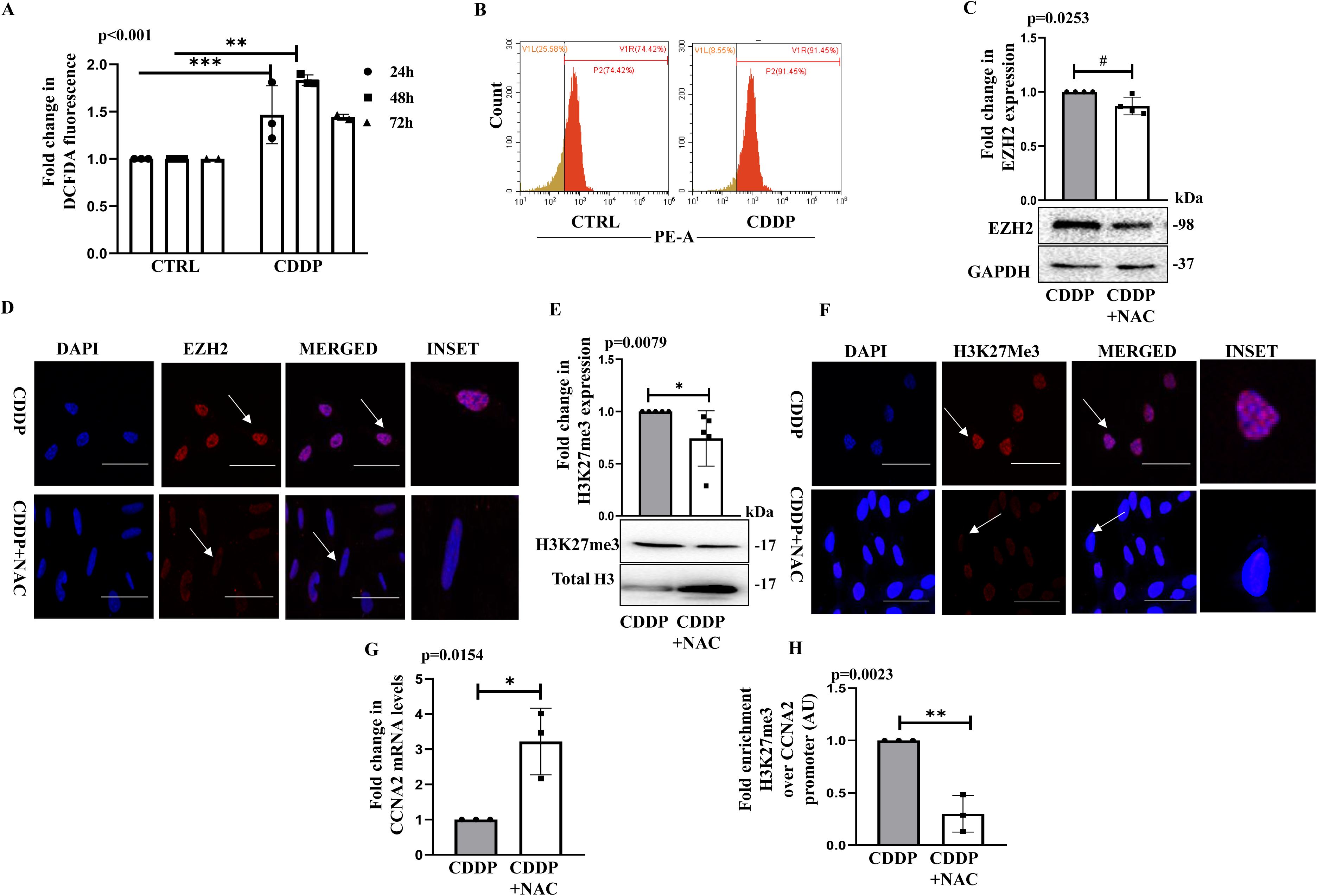
ROS regulates repressive epigenomic marks responsible for growth arrest. **(A)** Reactive Oxygen species levels after incubating cells with CDDP for various time points (24h, 48h, 72h) as measured through DCFDA staining. **(B)** Flow cytometric analysis of mitochondrial ROS generation *via* MitoSox at 24h. **(C)** Immunoblot analysis showing expression of EZH2 after removal of oxidative stress (with NAC) at 24h; GAPDH is the loading control. **(D)** Immunofluorescence staining showing the expression of EZH2 at 24h (Scale Bar: 20 μm). **(E)** Immunoblot analysis showing expression of H3K27me3 after removal of oxidative stress (with NAC) at 24h; total H3 is the loading control. **(F)** Immunofluorescence staining showing the expression of H3K27me3 at 24h (Scale Bar: 20 μm). **(G)** Change in mRNA expression of cell cycle gene CCNA2 at 24h. **(H)** Enrichment of H3K27me3 over the cell cycle genes (CCNA2) as analyzed through ChIP-qPCR. All values are represented as mean±SD; n=3. Unpaired t-test and One-way ANOVA, wherever applicable, were applied with (*) p<0.05 to estimate the significance when two groups were compared. CTRL represents untreated cells.

To investigate the role of ROS in mediating epigenetic alterations following CDDP treatment, we pre-treated cells with the ROS scavenger N-acetyl cysteine (NAC) prior to CDDP exposure. Notably, ROS quenching resulted in a marked reduction in EZH2 protein levels, as shown by immunoblotting **(Fig 3C; Supplementary Figure 2B)**. While NAC treatment did not alter the subcellular localization of EZH2, a significant decrease in its intranuclear fluorescence intensity was observed **(Fig 3D; Supplementary Figure 2C)**. Consistent with the reduction in EZH2 expression, a corresponding decrease in the levels of its histone substrate, H3K27me3, was detected by both immunoblotting and immunofluorescence **(Fig 3E, 3F; Supplementary Figure 2D–E)**. Given the earlier observation of reduced CCNA2 expression following CDDP treatment, we next examined whether this effect was ROS-dependent. Strikingly, NAC co-treatment restored CCNA2 expression and this was accompanied by a reduced enrichment of H3K27me3 at its promoter region **(Figures 3G and 3H)**, suggesting a probable re-induction of the cellular proliferative ability **(Fig 3G and 3H**). Collectively, these findings indicate that ROS acts as a critical mediator of specific epigenetic modifications induced by CDDP, particularly through modulation of EZH2 and associated H3K27me3, ultimately contributing to the regulation of cellular proliferation.

### 3.4. YAP is activated post-CDDP exposure-a result of ROS

In quest of key molecules that can regulate the ROS-mediated epigenetic changes post CDDP exposure, we further analyzed existing literature and RNA sequencing data to excavate the probable key modulators. Notably, recent literature, together with RNA sequencing analyses of CDDP-treated HOS cells, revealed compelling evidence of dysregulation in multiple genes associated with the regulation of cellular cytoskeletal dynamics (GO:0015630) (14, 25, 26). One such aberrantly expressed pathway was the Hippo Signaling Pathway, and therefore, we further explored its probable role in the regulation of ROS-mediated epigenome under CDDP stress. The Hippo signaling pathway is well-established in regulating contact inhibition and cellular mechano-sensing (27, 28). In line with its tumor-suppressive function, we observed that LATS1-the core kinase of the Hippo pathway responsible for inhibiting the transcriptional co-activator Yes-Associated Protein (YAP)—was significantly downregulated following CDDP exposure (LogL fold change = –3.39; *p* = 0.022; GEO Accession: GSE86053). This indicates a probable activation of YAP. Immunohistochemistry-based analyses have shown that high YAP expression is observed in approximately 60–80% of OS patients and is frequently associated with poor prognosis (29). Importantly, in our study, low-dose CDDP treatment resulted in a transcriptional upregulation of YAP and its downstream target genes, including CYR61 and CTGF **(Figure 4A; Supplementary Figures 3A–B)**. We further observed a significant increase in YAP protein levels following CDDP treatment, which was accompanied by a concomitant decrease in the expression of its upstream negative regulator, LATS1 (**Fig 4B and Supplementary Figure 3C**). Consistent with our earlier findings, ROS quenching resulted in a reduction of YAP expression at both the mRNA and protein levels **(Fig 4C and 4D)**, and led to reduced nuclear localization of YAP **(Fig 4E)**. In addition, a significant decrease in the mRNA of downstream effectors of YAP, and a reversion in expression of LATS1 was also observed with NAC **(Supplementary Figure 3D-F)**. To further validate these findings, we conducted a luciferase reporter assay using a YAP-responsive promoter construct. Consistent with the observed increases in YAP mRNA and protein levels, CDDP treatment significantly enhanced YAP transcriptional activity, as indicated by increased luciferase signal. In contrast, co-treatment with NAC markedly reduced luciferase activity **(Fig 4F; Supplementary Figure S3G)**.

**Figure 4.**
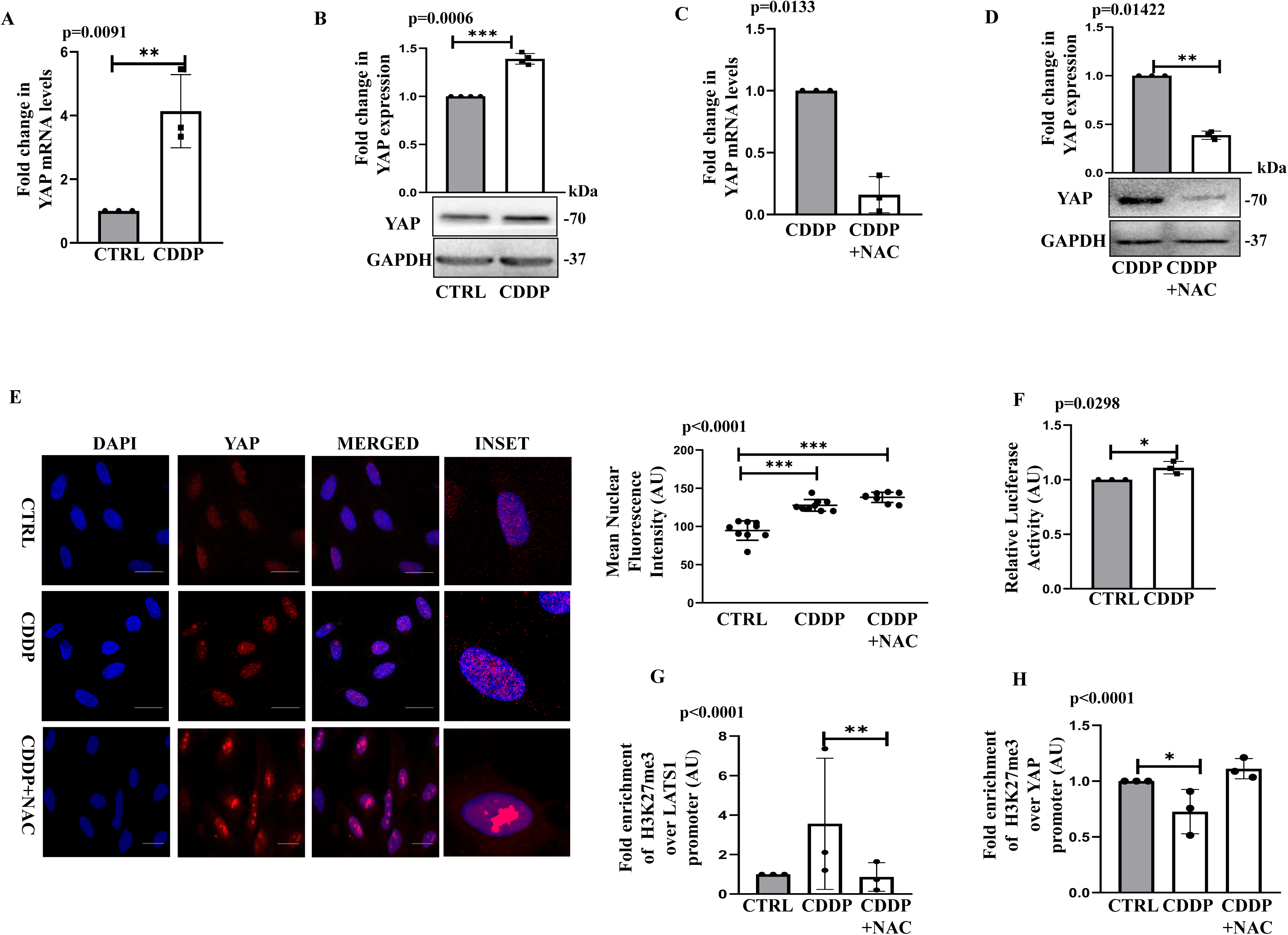
YAP is activated post-CDDP exposure-a result of ROS. **(A)** Change in mRNA expression of YAP at 24h when exposed to a sub-lethal dose of CDDP. **(B)** Immunoblot analysis showing expression of YAP at 24h, GAPDH is the loading control. **(C)** Change in mRNA expression of YAP at 24h after ablation of oxidative stress. **(D)** Immunoblot analysis showing expression of YAP at 24h, GAPDH is the loading control. **(E)** Immunofluorescence staining showing the expression of YAP at 24h (Scale Bar: 20 μm). **(F)** Change in luciferase activity of YAP-responsive promoter construct post exposure to drug treatment at 24h. **(G, H)** Enrichment of H3K27me3 over the Hippo pathway core genes-LATS1 and YAP, respectively, through ChIP-qPCR. All values are represented as mean±SD; n=3. *, ** and *** refers to p value significance of ≤0.01, ≤0.001 & ≤0.0001 respectively.

Collectively, these data establish a link between CDDP-induced oxidative stress and enhanced YAP activity. However, given the overall increase in repressive histone modifications following CDDP treatment, we sought to investigate the underlying mechanism driving the upregulation of YAP. Interestingly, ChIP-qPCR analysis targeting H3K27me3 revealed increased enrichment of this repressive histone mark at the promoter regions of LATS1 following CDDP exposure **(Fig 4G)**. In contrast, ROS quenching with NAC reversed this enrichment pattern, suggesting that CDDP suppresses LATS1 transcription through ROS-mediated epigenetic silencing, thereby promoting YAP stabilization and activation. Notably, an opposing enrichment profile was observed at the YAP promoter, where H3K27me3 levels were reduced in response to CDDP **(Fig 4H)**, indicating that YAP may also be subject to independent epigenetic regulation under drug-induced stress. Herein, the overall findings suggest that CDDP-induced oxidative stress results in an increased YAP activity; we hence thereafter sought to investigate whether YAP has any role in the regulation of the repressive epigenomic marks under CDDP stress.

### 3.5. Pharmacological/genetic ablation of YAP decreases repressive chromatin marks

To elucidate the role of YAP, we employed the well-characterized pharmacological inhibitor Verteporfin (VP) and also conducted genetic knockdown studies using siRNA. The inhibitory concentration of VP was initially evaluated through MTT assay, and through monitoring the expression of YAP in the presence of VP (10 uM) **(Supplementary Fig4A and 4B)**. Notably, pharmacological inhibition of YAP attenuated the CDDP-induced elevation of global H3K27me3 levels. **(Fig 5A)**. Similar results were also obtained upon siRNA-mediated knockdown of YAP **(Fig 5B and Supplementary Fig 4C)**. Collectively, these observations support the hypothesis that YAP plays a critical role in mediating the induction of global and or specific repressive histone modifications in response to drug-induced stress. Moreover, YAP expression exhibited a positive association with EZH2 expression. Importantly, VP exposure or siRNA-mediated knockdown of YAP resulted in reduced EZH2 transcript and protein levels **(Fig 5C and 5D**). Furthermore, EZH2 nuclear localization was markedly reduced following CDDP and VP or siYAP treatment compared to CDDP alone **(Fig 5E)**, suggesting that YAP regulates not only the deposition of repressive methylation marks but also the expression and sub-cellular localization of the methyltransferase – EZH2. We additionally conducted co-localization studies of YAP and EZH2 following CDDP treatment, and notably, immunofluorescence and co-immunoprecipitation analyses demonstrated a physical interaction between the two proteins post-CDDP exposure (**Fig 5F and 5G)**. The abovementioned findings provide clear evidence that YAP regulates repressive epigenetic marks in OS cells by interacting with and modulating the expression of EZH2 following drug-induced stress. Importantly, pharmacological inhibition of EZH2 using GSK126 did not significantly affect YAP levels **(Supplementary Fig 4D)**. Collectively, these datasets reinforce the pivotal role of Hippo/YAP signaling in mediating epigenomic alterations in OS cells post drug exposure.

**Figure 5.**
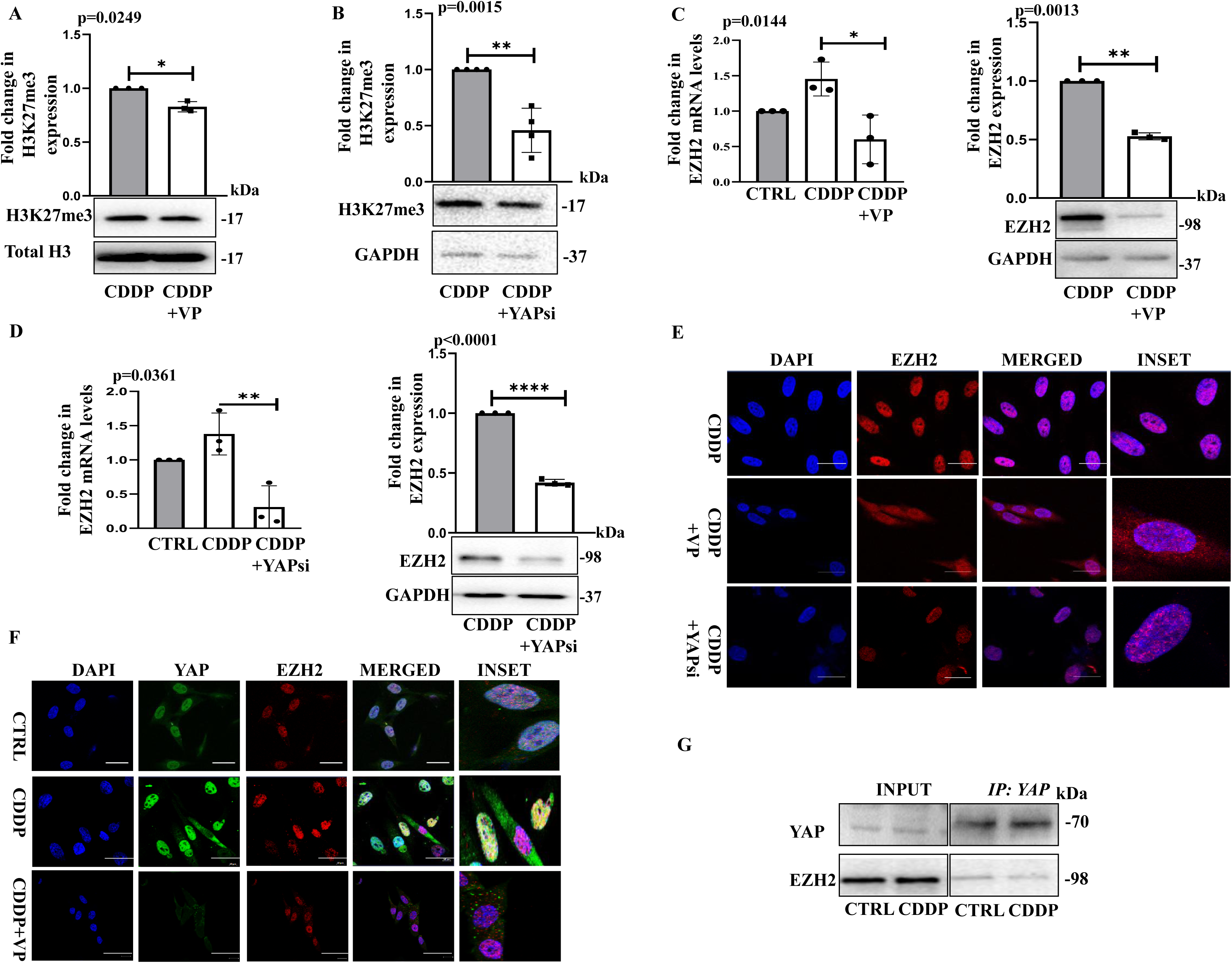
Pharmacological/genetic ablation of YAP decreases the repressive chromatin mark. **(A)** Change in H3K27me3 expression through immunoblot post CDDP plus VP (10 μM) treatment. **(B)** Immunoblot analysis showing expression of H3K27me3 after siRNA-mediated knockdown of YAP 24h post CDDP exposure; GAPDH is the loading control. **(C)** Change in mRNA and protein expression of EZH2 at 24h when YAP is inhibited with VP (10 µM) or **(D)** after siRNA treatment (40 nm), respectively; GAPDH is the loading control. **(E)** Immunofluorescence staining showing the expression of EZH2 at 24h (Scale Bar: 20 μm) when YAP is ablated *via* VP or siRNA. **(F)** Immunofluorescence staining showing the expression and co-localization of YAP and EZH2 at 24h with CDDP, and CDDP plus VP (10 µM) (Scale Bar: 20 μm). **(G)** Immunoblot analysis showing physical interaction between EZH2 and YAP at 24h, after CDDP exposure. All values are represented as mean±SD; n=3. *, ** and *** refers to p value significance of ≤0.01, ≤0.001 & ≤0.0001 respectively.

### 3.6. Inhibition of YAP as a potent strategy to sensitize OS cells to CDDP

Given the increasing clinical adoption of combination therapies in cancer treatment, our findings offer a potential strategy to optimize OS therapy by enhancing the efficacy of low-dose CDDP. Based on the data presented above, we investigated whether pharmacological inhibition of YAP could sensitize OS cells to CDDP. Specifically, we evaluated the effect of Verteporfin (VP, 10 µM), an FDA-approved drug, in combination with low-dose CDDP. MTT assay results demonstrated that VP significantly enhanced the cytotoxic response to CDDP (3µM) (**Fig 6A**). This observation was further corroborated by Annexin V/PI staining, which revealed a marked increase in apoptotic Annexin V-positive cells following the combination treatment **(Fig 6B)**. Importantly, YAP inhibition by VP exerted greater cytotoxicity than EZH2 inhibition by GSK126 (25 µM) in the OS cells tested **(Fig 6C and 6D)**. These findings highlight the potential of combining low-dose CDDP with VP as an effective therapeutic approach, potentially reducing the adverse effects associated with high-dose chemotherapy. Nevertheless, *in vivo* investigations are required to comprehensively validate the therapeutic efficacy and safety profile of this combination strategy. Alternatively, inhibition of the histone deacetylases (SAHA, a pan-HDAC inhibitor), which remove histone acetyl groups to induce transcriptomic repression, also showed a synergy with CDDP, lowering CDDP-induced H3K27me3, and EZH2 **(Fig 6E and F)** and inducing increased cell death in combination **(Fig 6G and H)** suggesting that a reversal of repressive tags can also be explored as a therapeutic modality alongside CDDP.

**Figure 6.**
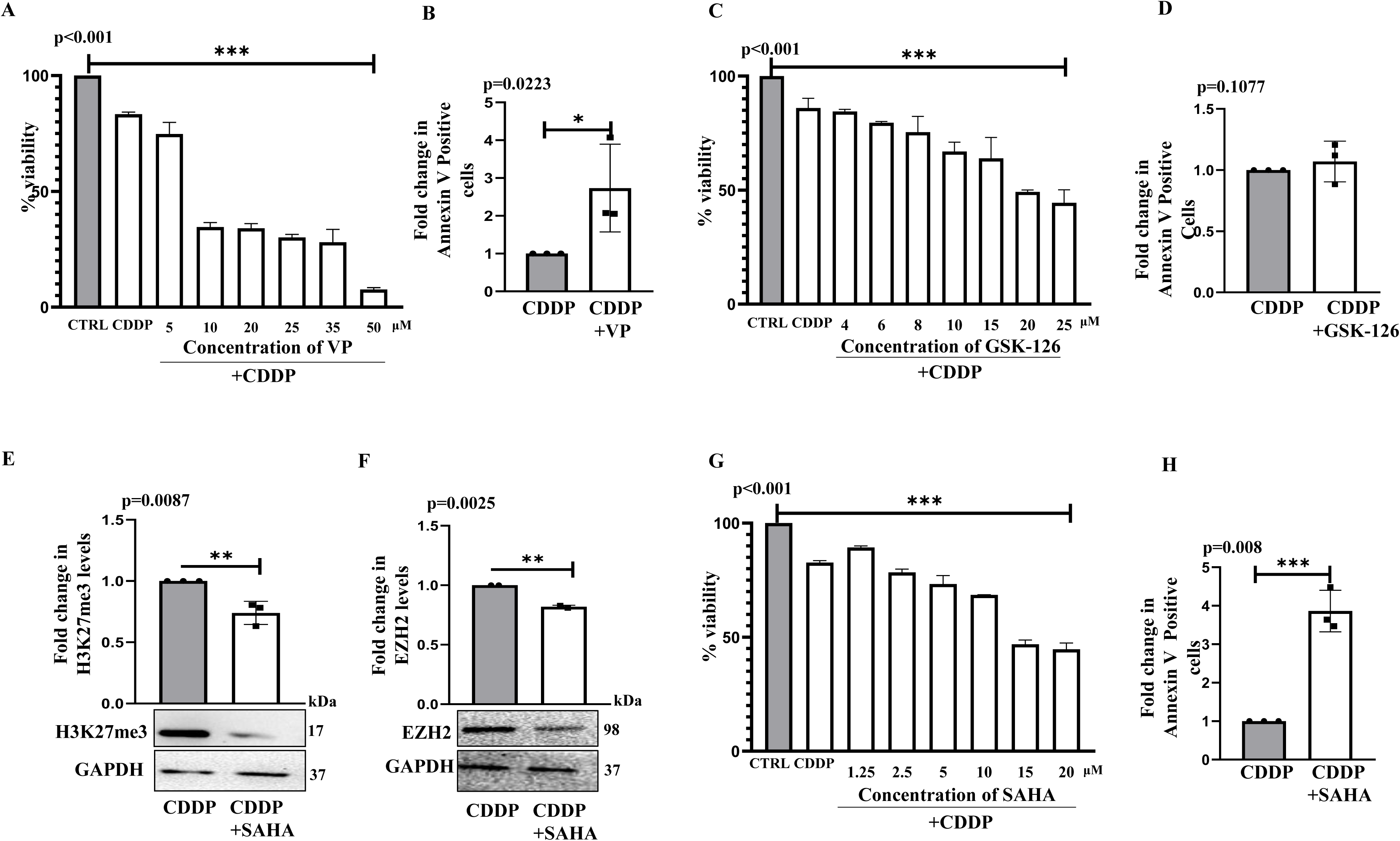
Inhibition of YAP and/or EZH2 as a potent strategy to sensitize OS cells. (A) Cell Viability (MTT) assay at different concentrations of VP and CDDP (3 µM) for 24h. (B) The bar graph illustrates the number of Annexin V-positive cells in CDDP plus VP (10 µM), compared to only CDDP-treated cells. (C) Cell Viability (MTT) assay at different concentrations of GSK-126 and CDDP (3 µM) for 24h. (D) The bar graph illustrates the number of Annexin-V positive cells in CDDP plus GSK-126 (25 µM), compared to only CDDP-treated cells. (E) Change in H3K27me3 expression through immunoblot post CDDP plus SAHA (15 µM) treatment. **(F)** Immunoblot analysis showing expression of EZH2 after combined treatment of CDDP (3 µM) plus SAHA (15 µM) at 24h; GAPDH is the loading control. **(G)** Cell Viability (MTT) assay at different concentrations of SAHA and CDDP (3 µM) for 24h. **(H)** Bar graph illustrating the number of Annexin-V positive cells in CDDP plus SAHA (15 µM), compared to only CDDP-treated cells. All values are represented as mean±SD; n=3. *, **, ***refers to p value significance of ≤0.01, ≤0.001 & ≤0.0001 respectively. CTRL represents the untreated cells.

## 4. Discussion

Cisplatin has long served as a first-line chemotherapeutic agent across various malignancies. However, its clinical efficacy is often limited by the development of intrinsic or acquired resistance. While platinum-based drug re-treatment strategies have gained attention, a clear understanding of optimal combination regimens and their applicability across different cancer types remains elusive. Several meta-analyses have supported the use of low-dose cisplatin regimens, citing their potent anti-tumor activity and potential utility as adjuvant therapies or in combination with other molecular modulators (30–32). Herein, in the context of osteosarcoma, there still remains a lack of effective and standardized therapeutic regimens, with current treatment approaches relying primarily on conventional chemotherapy. Therefore, comprehensive characterization of the molecular landscape of OS is essential for informing rational therapeutic strategies aimed at overcoming resistance and improving patient outcomes.

Herein, oxidative balance is essential for maintaining cellular homeostasis and influences the pathophysiology of connective tissues, such as bone (33). Emerging evidence indicates that certain tumor cells can reprogram their antioxidant defense mechanisms under the conditions of elevated oxidative stress, a process that fosters drug resistance (34, 35), particularly in advanced stages. Consequently, tumor cells adapted to sustained oxidative stress may exhibit reduced sensitivity to single-agent therapies, particularly platinum-based chemotherapeutics. Within this setting, oxidative stress is increasingly acknowledged as a key modulator of epigenomic reprogramming (5, 36). Therefore, disruption of redox homeostasis can affect the expression and activity of various chromatin-modifying enzymes, including histone deacetylases (e.g., HDAC1), histone methyltransferases (e.g., HMT1), histone acetyltransferases (e.g., HAT1), and DNA methylation-related enzymes such as DNMT1, DNMT3A, and MBD4 (37). Evidence suggests that ROS, can impair the DNA-binding capacity of DNA methyltransferases (DNMTs), leading to global DNA hypomethylation (38, 39). Furthermore, oxidative stress has been implicated in heterochromatin remodeling across various disease models, including cardiovascular pathologies (39). Importantly, oxidative stress influences both the histone substrates and their modifying enzymes. For example, JmjC domain-containing histone demethylases utilize Fe(II) and α-ketoglutarate (αKG) as essential cofactors to catalyze oxidative demethylation through hydroxymethyl lysine intermediates (37). Under conditions of oxidative stress, the functional integrity of these cofactors is compromised, resulting in increased histone methylation. Conversely, removal of oxidative stress restores cofactor activity, leading to a reduction in repressive marks such as H3K27me3. Our findings support this mechanism, demonstrating that even sub-toxic doses of cisplatin can induce repressive histone modifications, which are reversible upon ROS quenching. These observations highlight intracellular ROS as a potentially universal modulator of the epigenome, capable of regulating gene expression by influencing histone dynamics. Such global epigenetic alterations may contribute to therapeutic resistance by silencing tumor suppressor genes, reducing accessibility of DNA-damaging drugs, and/or activating oncogenes. Notably, chromatin-modifying enzymes are frequently mutated in cancers, where partial loss (haploinsufficiency) can drive tumor progression, while complete loss is often deleterious to cells. This characteristic makes epigenetic therapies, or “epi-drugs,” a promising strategy to overcome traditional resistance mechanisms and improve treatment efficacy.

As we explored key drivers of epigenetic modifiers, there were several molecular candidates that can potentially modulate the epigenomic players. For example, aberrant activation of the YAP/TAZ transcriptional co-activators has been reported across multiple solid malignancies, supported with substantial genetic evidence. Herein, previous studies have also reported that YAP can interact with the nucleosome remodelling and deacetylase (NuRD) complex as well to facilitate histone deacetylation at specific gene promoters, thereby enabling recruitment of the Polycomb Repressive Complex 2 (PRC2) and subsequent gene silencing. Conventionally, the Hippo-YAP signaling axis has been implicated in mediating drug resistance and tolerance, ultimately contributing to tumor relapse (40, 41). 11, 42, 43). In alignment with these findings, our study demonstrates that exposure to cisplatin leads to increased YAP activity. However, whether YAP exerts its oncogenic functions solely through canonical Hippo pathway signalling remained an open question. We therefore analyzed the potential role of YAP in modulating the epigenome as an additional mechanism to influence cellular behaviour and therapeutic response. In our study, we observed predominant nuclear localization of YAP, consistent with findings from both patient samples and murine models (41). Importantly, YAP knockdown led to a transcriptional downregulation of EZH2, the catalytic component of PRC2, indicating a functional link between YAP and EZH2 expression. We further demonstrated that YAP and EZH2 not only co-localize within the nucleus but also physically interact. Notably, silencing YAP resulted in a significant reduction in the nuclear localization of EZH2, suggesting that YAP may contribute to the nuclear stabilization or retention of EZH2 and its associated repressive functions. Downregulation of YAP, either through direct silencing or via ROS quenching, was found to significantly alter EZH2 expression patterns, indicating that YAP functions upstream of EZH2 in this regulatory cascade. Based on these observations, we propose that YAP promotes tumor cell adaptation to chemotherapeutic stress by orchestrating EZH2-mediated transcriptional repression of critical regulatory genes. Collectively, our findings identify a novel ROS–YAP–EZH2 oncogenic axis that can be an efficient way to molecularly nail drug resistance in OS patients. Moreover, our results support the rationale for incorporating epigenetic modulators (epi-drugs) to sensitize tumor cells to conventional chemotherapeutics, thereby enabling the use of lower, less toxic doses and potentially improving patient outcomes.

## 5. Data Availability Statement

All original data generated in the study are provided within the article and its supplementary material. Requests for additional information should be directed to the corresponding author(s).

## 6. Funding

This study was supported by DST-SERB (CRG/2022/005172) of SM and ICMR of RC (2020–1404/ADHOC-BMS).

## Acknowledgements

AD gratefully acknowledges the Council of Scientific and Industrial Research (CSIR), India, for the award of a research fellowship. The authors also acknowledge the infrastructural support provided by BITS Pilani, Pilani Campus

## 7. Conflict of Interest

The authors declare that there are no conflicts of interest.

## Supplementary Figure Legends

**Supplementary Figure 1.** Change in mRNA expression of HDAC5 **(A)**, HDAC6 **(B)**, HDAC9 **(C)** post low-dose CDDP exposure (3 µM, 24h). All values are represented as mean±SD; n=3. *, ** and *** refers to p value significance of ≤0.01, ≤0.001 & ≤0.0001 respectively.

**Supplementary Figure 2. (A)** Mitochondrial membrane potential post CDDP treatment; shift in red to green fluorescence in JC-1 dye as analyzed through flow cytometry. **(B)** Immunoblot showing expression of EZH2 after CDDP plus NAC treatment at 24h, untreated cells are used as a control. **(C)** Immunofluorescence staining showing the expression of EZH2 post CDDP plus NAC treatment at 24h (Scale Bar: 20 μm). (**D)** Immunoblot showing expression of H3K27me3 post CDDP plus NAC treatment at 24h, untreated cells are used as a control. **(E)** Immunofluorescence staining showing the expression of H3K27me3 post CDDP plus NAC treatment at 24h (Scale Bar: 20 μm). All values are represented as mean±SD; n=3. *, ** and *** refers to p value significance of ≤0.01, ≤0.001 & ≤0.0001 respectively.

**Supplementary Figure 3. (A)** Change in mRNA expression of CTGF, post CDDP treatment at 24h, untreated cells are used as control. **(B)** Change in mRNA expression of CYR61, post CDDP treatment at 24h, untreated cells are used as control. **(C)** Immunoblot showing the change in expression of LATS1, post-CDDP treatment at 24h. **(D)** Change in mRNA expression of CTGF post CDDP plus NAC treatment at 24h, untreated cells are used as control. **(E)** Change in mRNA expression of CYR61 post CDDP plus NAC treatment at 24h, untreated cells are use as control. **(F)** Immunoblot showing expression of LATS1 post CDDP plus NAC treatment at 24h, untreated cells are used as control. **(G)** Change in luciferase activity of YAP post exposure to CDDP plus NAC treatment at 24h; untreated cells are used as control. All values are represented as mean±SD; n=3. *, ** and *** refers to p value significance of ≤0.01, ≤0.001 & ≤0.0001 respectively.

**Supplementary Figure 4. (A)** Cell viability (MTT) assay at different concentrations of VP VC: Vehicle control, (DMSO, 0.1%). **(B)** Immunoblot showing expression of YAP post CDDP and VP (10 μM) at 24h, untreated cells are used as a control. **(C)** Immunoblot showing expression of YAP post siRNA knockdown of YAP at 24h, untreated cells are as control. **(D)** Immunoblot analysis showing expression of EZH2 and YAP at 24h post CDDP and GSK-126 (25 μM); GAPDH is used as the loading control. All values are represented as mean±SD; n=3. *, ** and *** refers to p value significance of ≤0.01, ≤0.001 & ≤0.0001 respectively.

## Notes

### Competing Interest Statement

The authors have declared no competing interest.

## References

1. Bielack SS, Kempf-Bielack B, Delling Gn, Exner GU, Flege S, Helmke K, et al. Prognostic factors in high-grade osteosarcoma of the extremities or trunk: an analysis of 1,702 patients treated on neoadjuvant cooperative osteosarcoma study group protocols. 2002;20(3):776–90. DOI: 10.1200/JCO.2002.20.3.776

2. Hamre MR, Severson RK, Chuba P, Lucas DR, Thomas RL, Mott MPJR, et al. Osteosarcoma as a second malignant neoplasm. 2002;65(3):153–7. DOI: 10.1016/s0167-8140(02)00150-0

3. Bielack SS, Carrle D, Hardes J, Schuck A, Paulussen MJCtoio. Bone tumors in adolescents and young adults. 2008;9:67–80. DOI: 10.1007/s11864-008-0057-1

4. Brown A, Kumar S, Tchounwou PBJJocs, therapy. Cisplatin-based chemotherapy of human cancers. 2019;11(4).

5. García-Guede Á, Vera O, Ibáñez-de-Caceres IJA. When oxidative stress meets epigenetics: implications in cancer development. 2020;9(6):468. DOI: 10.3390/antiox9060468

6. Mahalingaiah PKS, Ponnusamy L, Singh KPJO. Oxidative stress-induced epigenetic changes associated with malignant transformation of human kidney epithelial cells. 2017;8(7):11127. DOI: 10.18632/oncotarget.12091

7. Nishida N, Arizumi T, Takita M, Kitai S, Yada N, Hagiwara S, et al. Reactive oxygen species induce epigenetic instability through the formation of 8-hydroxydeoxyguanosine in human hepatocarcinogenesis. 2013;31(5-6):459–66. DOI: 10.1159/000355245

8. Hahm JY, Park J, Jang E-S, Chi SWJE, Medicine M. 8-Oxoguanine: From oxidative damage to epigenetic and epitranscriptional modification. 2022;54(10):1626–42. DOI: 10.1038/s12276-022-00822-z

9. Weitzman SA, Turk PW, Milkowski DH, Kozlowski KJPotNAoS. Free radical adducts induce alterations in DNA cytosine methylation. 1994;91(4):1261–4. DOI: 10.1073/pnas.91.4.1261

10. Zhao B, Tumaneng K, Guan K-LJNcb. The Hippo pathway in organ size control, tissue regeneration and stem cell self-renewal. 2011;13(8):877–83. DOI: 10.1038/ncb2303

11. Feng X, Degese MS, Iglesias-Bartolome R, Vaque JP, Molinolo AA, Rodrigues M, et al. Hippo-independent activation of YAP by the GNAQ uveal melanoma oncogene through a trio-regulated rho GTPase signaling circuitry. 2014;25(6):831–45. DOI: 10.1016/j.ccr.2014.04.016

12. Han H, Yang B, Nakaoka HJ, Yang J, Zhao Y, Le Nguyen K, et al. Hippo signaling dysfunction induces cancer cell addiction to YAP. 2018;37(50):6414–24. DOI: 10.1038/s41388-018-0419-5

13. Venkatasubramanian G, Kelkar DA, Mandal S, Jolly MK, Kulkarni MJJoCM. Analysis of yes-associated protein-1 (YAP) target gene signature to predict progressive breast cancer. 2022;11(7):1947. DOI: 10.3390/jcm11071947

14. Li S, Zhang X, Zhang R, Liang Z, Liao W, Du Z, et al. Hippo pathway contributes to cisplatin resistant-induced EMT in nasopharyngeal carcinoma cells. 2017;16(17):1601–10. DOI: 10.1080/15384101.2017.1356508

15. Zucchini C, Manara MC, Cristalli C, Carrabotta M, Greco S, Pinca RS, et al. ROCK2 deprivation leads to the inhibition of tumor growth and metastatic potential in osteosarcoma cells through the modulation of YAP activity. 2019;38:1–14. DOI: 10.1186/s13046-019-1506-3

16. Wu T, Hu H, Zhang T, Jiang L, Li X, Liu S, et al. miR-25 promotes cell proliferation, migration, and invasion of non-small-cell lung cancer by targeting the LATS2/YAP signaling pathway. 2019;2019. DOI: 10.1155/2019/9719723

17. Shao D, Zhai P, Del Re DP, Sciarretta S, Yabuta N, Nojima H, et al. A functional interaction between Hippo-YAP signalling and FoxO1 mediates the oxidative stress response. 2014;5(1):3315. DOI: 10.1038/ncomms4315

18. Venkataramani V, Küffer S, Cheung KC, Jiang X, Trümper L, Wulf GG, et al. CD31 expression determines redox status and chemoresistance in human angiosarcomas. 2018;24(2):460–73. DOI: 10.1158/1078-0432.CCR-17-1778

19. Hillmer RE, Link BAJC. The roles of Hippo signaling transducers Yap and Taz in chromatin remodeling. 2019;8(5):502. DOI: 10.3390/cells8050502

20. Skibinski A, Breindel JL, Prat A, Galván P, Smith E, Rolfs A, et al. The Hippo transducer TAZ interacts with the SWI/SNF complex to regulate breast epithelial lineage commitment. 2014;6(6):1059–72. DOI: 10.1016/j.celrep.2014.02.038

21. Ayyagari VN, Hsieh T-hJ, Diaz-Sylvester PL, Brard LJBc. Evaluation of the cytotoxicity of the Bithionol-cisplatin combination in a panel of human ovarian cancer cell lines. 2017;17(1):1–15. DOI: 10.1186/s12885-016-3034-2

22. Sharma SV, Lee DY, Li B, Quinlan MP, Takahashi F, Maheswaran S, et al. A chromatin-mediated reversible drug-tolerant state in cancer cell subpopulations. 2010;141(1):69–80. DOI: 10.1016/j.cell.2010.02.027

23. Liou G-Y, Storz PJFrr. Reactive oxygen species in cancer. 2010;44(5):479–96. DOI: 10.3109/10715761003667554

24. García-Giménez J-L, Garcés C, Romá-Mateo C, Pallardó FVJFRB, Medicine. Oxidative stress-mediated alterations in histone post-translational modifications. 2021;170:6–18. DOI: 10.1016/j.freeradbiomed.2021.02.027

25. Shimizu T, Fujii T, Sakai HJFic, biology d. The relationship between actin cytoskeleton and membrane transporters in cisplatin resistance of cancer cells. 2020;8:597835. DOI: 10.3389/fcell.2020.597835

26. Parker AL, Teo WS, McCarroll JA, Kavallaris MJIjoms. An emerging role for tubulin isotypes in modulating cancer biology and chemotherapy resistance. 2017;18(7):1434. DOI: 10.3390/ijms18071434

27. Gumbiner BM, Kim N-GJJocs. The Hippo-YAP signaling pathway and contact inhibition of growth. 2014;127(4):709–17. DOI: 10.1242/jcs.140103

28. Chang Y-C, Wu J-W, Wang C-W, Jang AC-CJFimb. Hippo signaling-mediated mechanotransduction in cell movement and cancer metastasis. 2020;6:157. DOI: 10.3389/fmolb.2019.00157

29. Zhan F, He T, Chen Z, Zuo Q, Wang Y, Li Q, et al. RhoA enhances osteosarcoma resistance to MPPa-PDT via the Hippo/YAP signaling pathway. 2021;11(1):1–19. DOI: 10.1186/s13578-021-00690-6

30. Xie X, Wu Y, Luo S, Yang H, Li L, Zhou S, et al. Efficacy and toxicity of low-dose versus conventional-dose chemotherapy for malignant tumors: a meta-analysis of 6 randomized controlled trials. 2017;18(2):479. DOI: 10.22034/APJCP.2017.18.2.479

31. Simsek C, Esin E, Yalcin SJJoo. Metronomic chemotherapy: a systematic review of the literature and clinical experience. 2019;2019. DOI: 10.1155/2019/5483791

32. Bocci G, Nicolaou K, Kerbel RSJCr. Protracted low-dose effects on human endothelial cell proliferation and survival in vitro reveal a selective antiangiogenic window for various chemotherapeutic drugs. 2002;62(23):6938–43.

33. Domazetovic V, Marcucci G, Iantomasi T, Brandi ML, Vincenzini MTJCCiM, Metabolism B. Oxidative stress in bone remodeling: role of antioxidants. 2017;14(2):209. DOI: 10.11138/ccmbm/2017.14.1.209

34. Liu Y, Li Q, Zhou L, Xie N, Nice EC, Zhang H, et al. Cancer drug resistance: redox resetting renders a way. 2016;7(27):42740. DOI: 10.18632/oncotarget.8600

35. Forman HJ, Zhang HJNRDD. Targeting oxidative stress in disease: Promise and limitations of antioxidant therapy. 2021;20(9):689–709. DOI: 10.1038/s41573-021-00233-1

36. Hernández-Cruz EY, Arancibia-Hernández YL, Loyola-Mondragón DY, Pedraza-Chaverri JJO. Oxidative Stress and Its Role in Cd-Induced Epigenetic Modifications: Use of Antioxidants as a Possible Preventive Strategy. 2022;2(2):177–210. 10.3390/oxygen2020015

37. Niu Y, DesMarais TL, Tong Z, Yao Y, Costa MJFRB, Medicine. Oxidative stress alters global histone modification and DNA methylation. 2015;82:22–8. DOI: 10.1016/j.freeradbiomed.2015.01.028

38. Yi X, Zhu Q-X, Wu X-L, Tan T-T, Jiang X-JJOM, Longevity C. Histone methylation and oxidative stress in cardiovascular diseases. 2022;2022. DOI: 10.1155/2022/6023710

39. Kietzmann T, Petry A, Shvetsova A, Gerhold JM, Görlach AJBJoP. The epigenetic landscape related to reactive oxygen species formation in the cardiovascular system. 2017;174(12):1533–54. DOI: 10.1111/bph.13792

40. Nguyen CD, Yi CJTic. YAP/TAZ signaling and resistance to cancer therapy. 2019;5(5):283–96. DOI: 10.1016/j.trecan.2019.02.010

41. Zanconato F, Cordenonsi M, Piccolo SJCc. YAP/TAZ at the roots of cancer. 2016;29(6):783–803. DOI: 10.1016/j.ccell.2016.05.005

42. Zhang Y, Li B, Shen L, Shen Y, Chen XJIjoi, pharmacology. The role and clinical significance of YES-associated protein 1 in human osteosarcoma. 2013;26(1):157–67. DOI: 10.1177/039463201302600115

43. Nguyen LT, Tretiakova MS, Silvis MR, Lucas J, Klezovitch O, Coleman I, et al. ERG activates the YAP transcriptional program and induces the development of age-related prostate tumors. 2015;27(6):797–808. DOI: 10.1016/j.ccell.2015.05.005

44. Saini H, Sharma H, Mukherjee S, Chowdhury S, Chowdhury RJCCI. Verteporfin disrupts multiple steps of autophagy and regulates p53 to sensitize osteosarcoma cells. 2021;21(1):1–16. DOI: 10.1186/s12935-020-01720-y

45. Pfaffl MWJNar. A new mathematical model for relative quantification in real-time RT–PCR. 2001;29(9):e45–e. DOI: 10.1093/nar/29.9.e45

