## Supplementary Figures for "Cisplatin-induced oxidative stress regulates YAP to modulate epigenome promoting survival of osteosarcoma cells"

**Supplementary Figure 1**

A

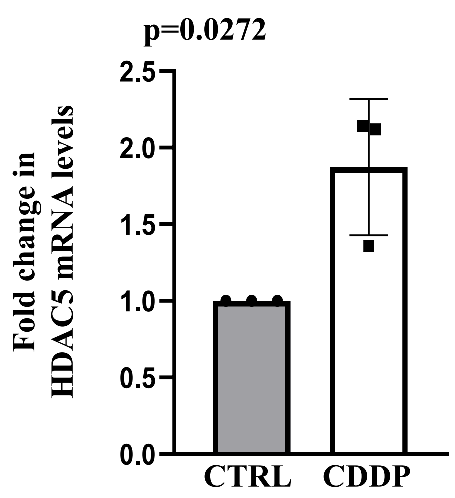


B

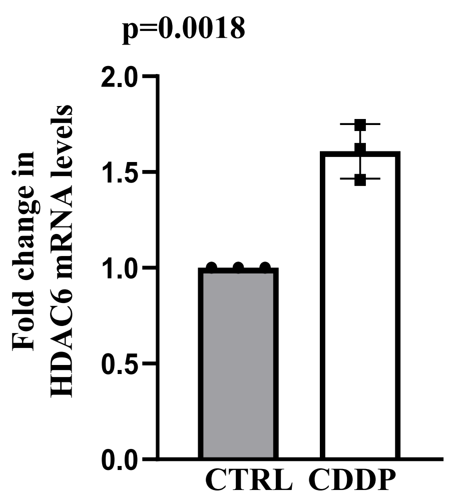


C

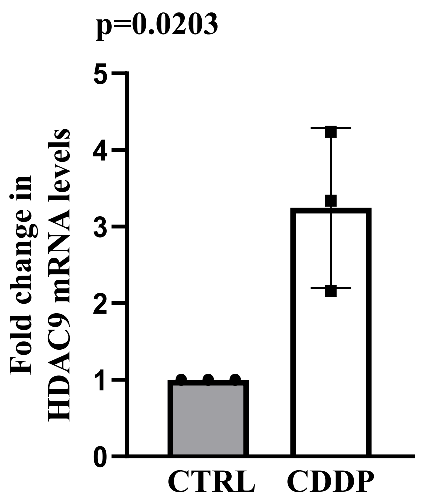


D

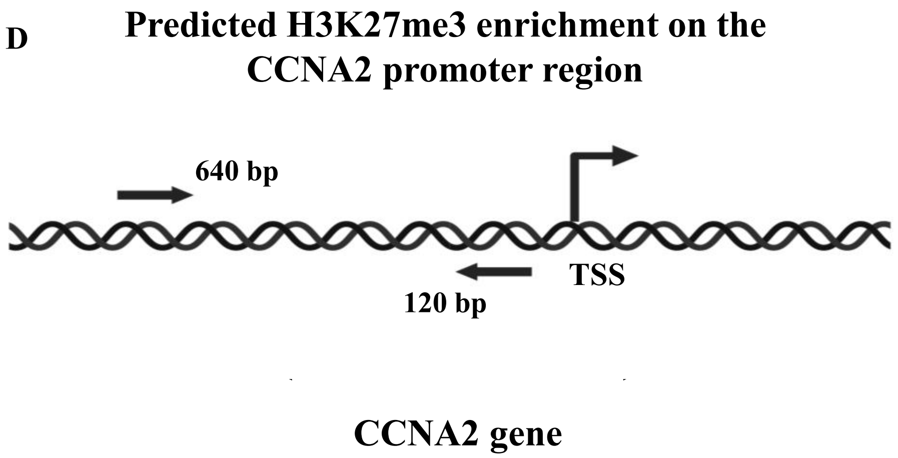

### Supplementary Figure 2

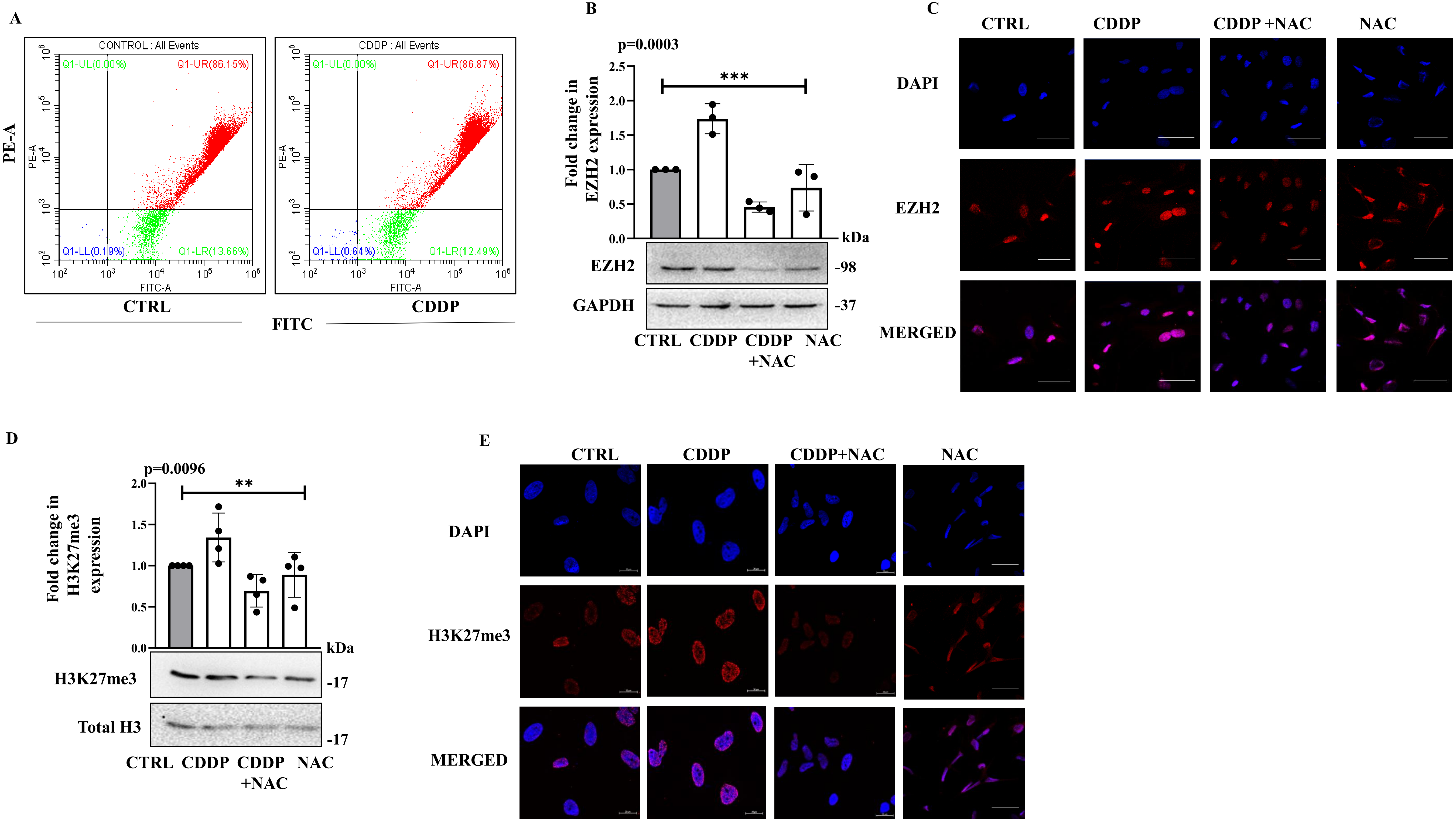

**Supplementary Figure 3**

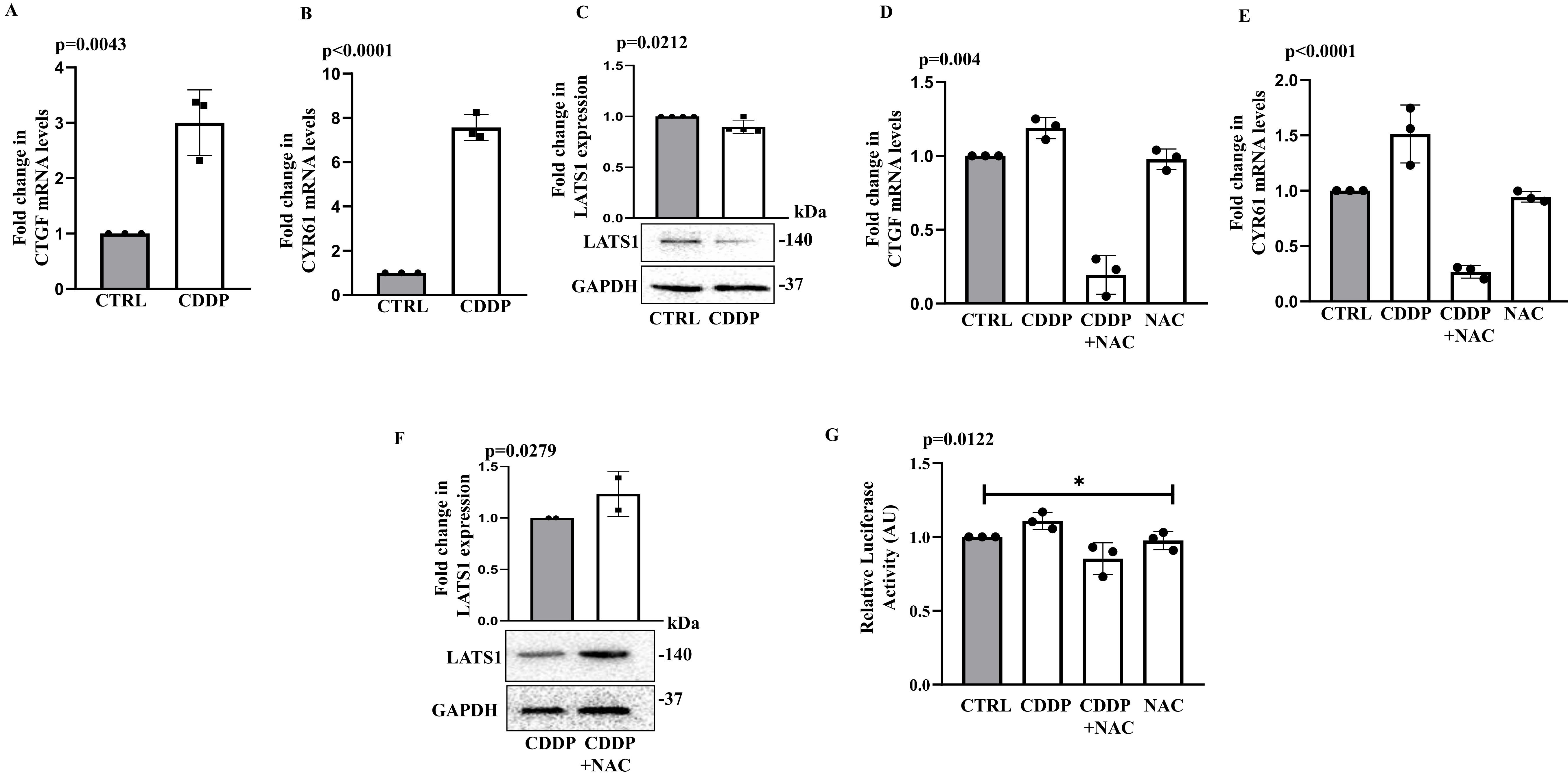

**Supplementary Figure 4**

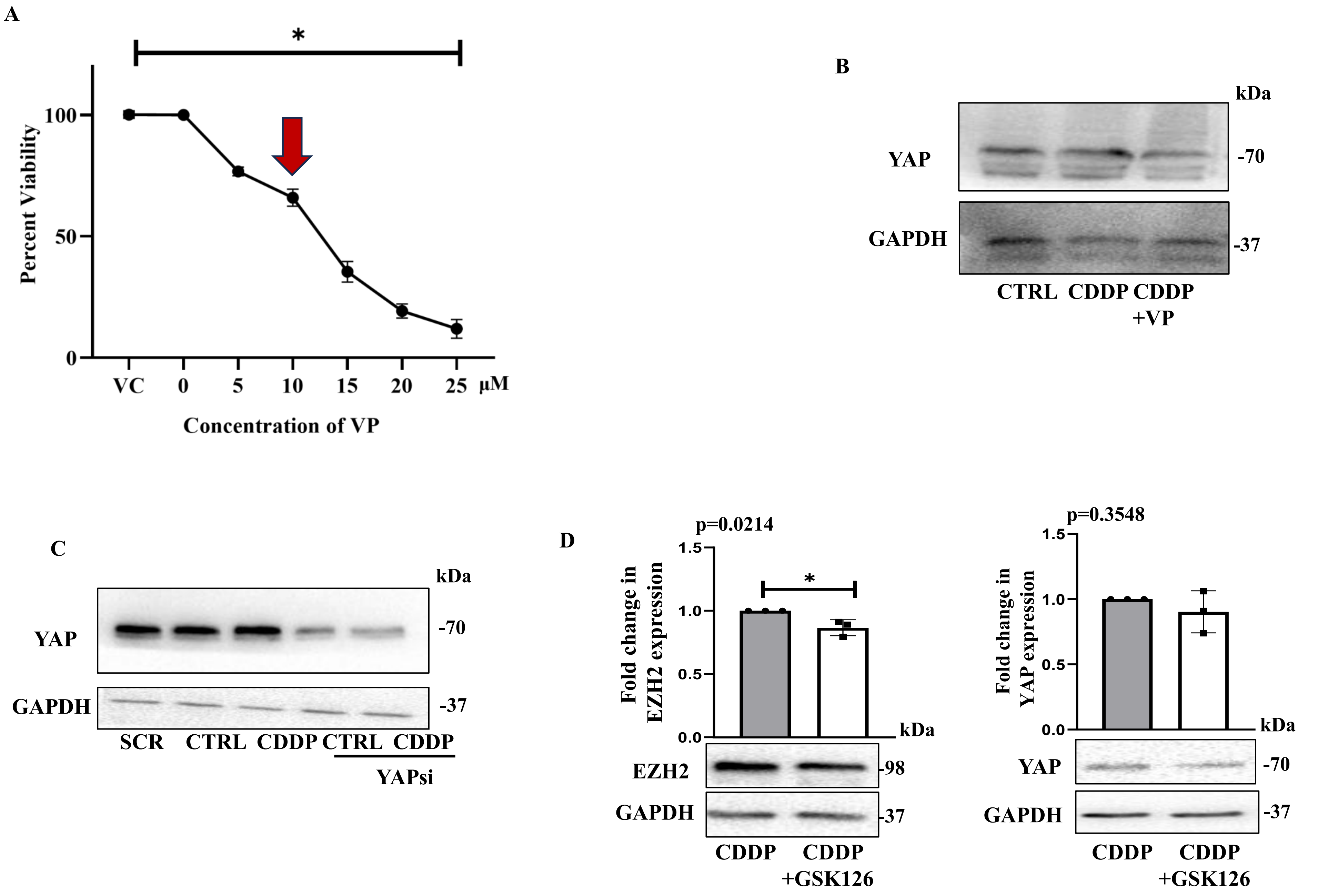
